# A phylogenomic approach reveals a low somatic mutation rate in a long-lived plant

**DOI:** 10.1101/727982

**Authors:** Adam J. Orr, Amanda Padovan, David Kainer, Carsten Külheim, Lindell Bromham, Carlos Bustos-Segura, William Foley, Tonya Haff, Ji-Fan Hsieh, Alejandro Morales-Suarez, Reed A. Cartwright, Robert Lanfear

## Abstract

Somatic mutations can have important effects on the life history, ecology, and evolution of plants, but the rate at which they accumulate is poorly understood, and has been very difficult to measure directly. Here, we demonstrate a novel method to measure somatic mutations in individual plants and use this approach to estimate the somatic mutation rate in a large, long-lived, phenotypically mosaic *Eucalyptus melliodora* tree. Despite being 100 times larger than *Arabidopsis*, this tree has a per-generation mutation rate only ten times greater, which suggests that this species may have evolved mechanisms to reduce the mutation rate per unit of growth. This adds to a growing body of evidence that illuminates the correlated evolutionary shifts in mutation rate and life history in plants.

---

Trees grow from multiple meristems which contain stem cells that divide to produce the somatic and reproductive tissues. A mutation occurring in a meristematic cell will be passed on to all resulting tissues, potentially causing an entire branch including leaves, stems, flowers, seeds, and pollen to have a genotype different from the rest of the plant^1,2^. These different genotypes may lead to phenotypic changes, potentially with important consequences for plant ecology and evolution^3-8^. For example, somatic mutations could explain how long-lived plants adapt to changing ecological conditions^9^, and are thought to influence long-term variation in rates of evolution and speciation among plant lineages^10^. Somatic mutations can degrade genetic stocks used in agriculture and forestry^11,12^, confer herbicide resistance to weed species^13^, and have been linked to declining plant fitness in polluted areas^14^. However, despite their importance and recent progress^1,2,15-18^, there remain significant analytical challenges in inferring somatic mutation rates from sequencing data in plants.

We present a novel solution to the challenges of measuring somatic mutation rate, leveraging the phylogeny-like structure of the tree itself to estimate the genome-wide somatic mutation rate in an individual plant by sequencing the full genome of three terminal leaves of eight branch tips from a single individual plant. Our strategy has three key advantages. First, using three biological replicates per branch tip significantly reduces the false-positive rate, because many types of error (e.g. sequencing error, or mutations induced during DNA extraction or library preparation) are very unlikely to appear at the same position in all three replicates, making it easy to distinguish these errors from biological signal. Second, our strategy includes an inbuilt positive control, because we can ask whether the phylogenetic tree we reconstruct from a given set of putative somatic mutations reflects the known physical structure of the tree (i.e. whether phylogeny correctly reconstructs ontogeny, as is expected for plant development in most cases). Third, the approach allows us to estimate the false negative and the false positive rate of our inferences directly from the replicate samples (see below, and methods).

We applied this approach to a long-lived yellow box (*Eucalyptus melliodora*) tree, notable for its phenotypic mosaicism: a single large branch in this individual is resistant to defoliation by Christmas beetles (*Anoplognathus* spp Coleoptera: Scarabaeidae) due to stable differences in leaf chemistry and gene expression^19,20^. We selected eight branch tips that maximized the intervening physical branch length on the tree (Figure 1), reasoning that this would increase our power by maximizing the number of sampled cell divisions and thus somatic mutations. We performed independent DNA extractions from three leaves from each branch tip, prepared three independent libraries for Illumina sequencing, and sequenced each library to 10x coverage (assuming a roughly 500 Mbp genome size, as is commonly observed in *Eucalyptus* species ^21^) using 100bp paired-end sequencing on an Illumina HiSeq 2500. Quality control of the sequence data verified that each sample was sequenced to ∼10x coverage, and that each branch tip was therefore sequenced to ∼30x coverage.

**Figure 1.**
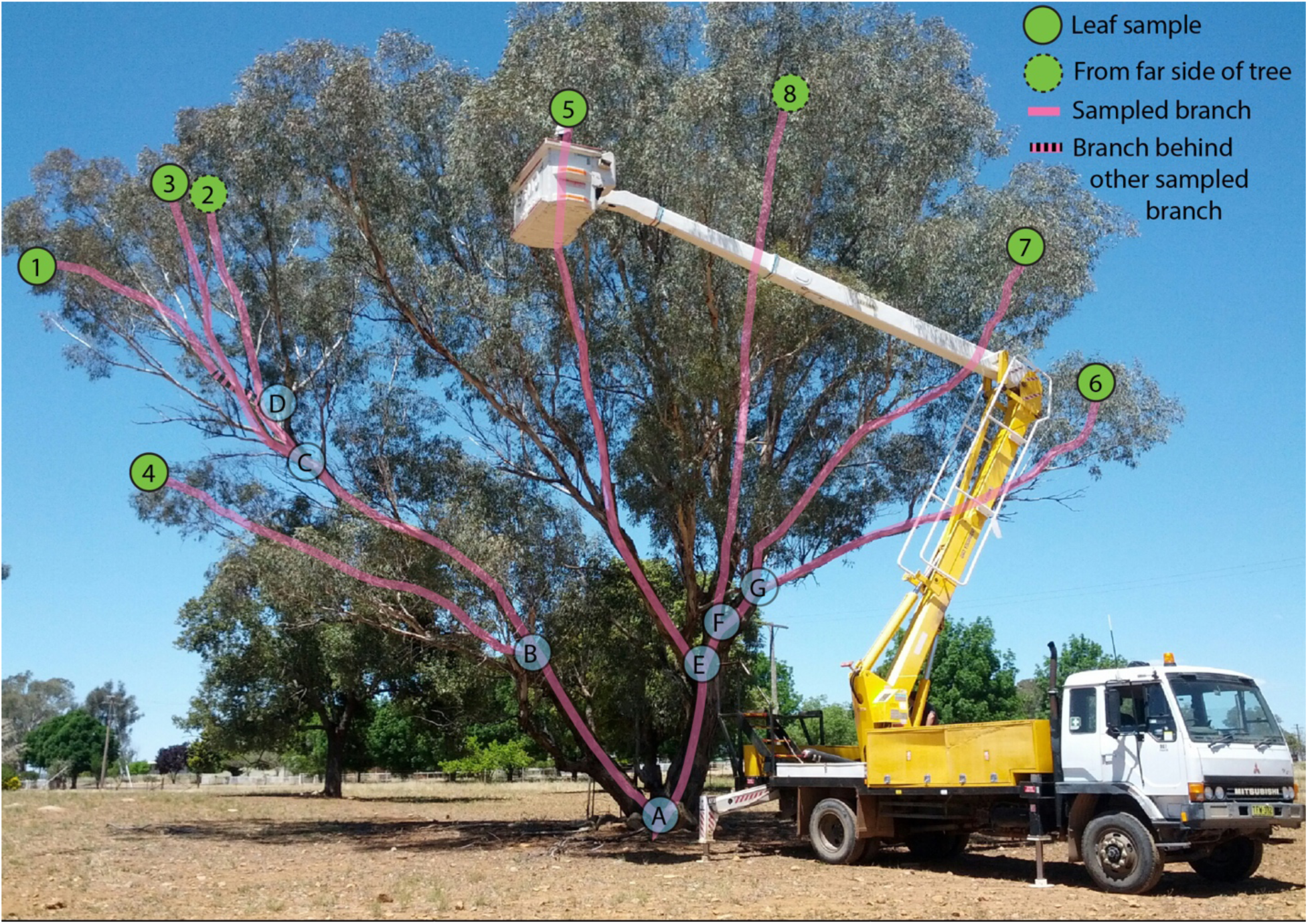
The *Eucalyptus melliodora* individual sequenced in this study. The eight branch tips sampled are shown by numbered green circles with internal nodes of the tree shown as letters in blue circles. Circles with dashed outlines are from the far side of the tree. Pink lines trace the physical branches that connect the sampled tips. The herbivore resistant branch comprises samples 1-3.

We first performed a positive control to confirm that the phylogeny of a set of high-confidence somatic variants matches the physical structure of the tree. To do this, we created a pseudo-reference genome by using our data to update the genome of *Eucalyptus grandis* (see methods). We then called variants using GATK^22^ in all three replicates, and used a set of strict filters to arrive at an alignment of 99 high-confidence somatic variants. To find the phylogenetic trees that best explain this alignment, we calculated the alignment’s parsimony score on all 10,395 possible phylogenetic trees of eight taxa. Parsimony is an appropriate method here because we do not expect more than one mutation to occur at any single site on any single branch of the *E. melliodora* tree. We selected the three phylogenetic trees with the most parsimonious scores and then asked whether these trees were more similar to the physical structure of the tree than would be expected by chance. To do this, we used the Path Difference to compare the structure of the physical tree to each of the three most parsimonious trees. We then compared these differences to the null distribution of Path Differences generated by comparing the structure of the physical tree to all possible 10,395 trees of eight taxa (Figure 2A). All three maximum parsimony trees were significantly more similar to the physical tree than would be expected by chance (p<0.001 in all cases, Figure 2A, dashed red lines). Furthermore, one of the most parsimonious trees is identical to the structure of the physical tree, and a maximum likelihood tree calculated from the same data shows just one topological difference compared to the structure of the physical tree, in which sample 8 is incorrectly placed as sister to sample 5, but with low bootstrap support of 44% (Figure 2A, blue line; Figure 2B). As would be expected if plants accumulate somatic mutations as they grow, there is a significant correlation between the branch lengths of the physical tree and the maximum-parsimony tree of the same topology (linear model forced through the origin: R^2^ = 0.82, p<0.001). These analyses demonstrate that the phylogeny recovered from the genomic data matches the physical structure of the tree, and confirm that there is strong biological signal in our data.

**Figure 2.**
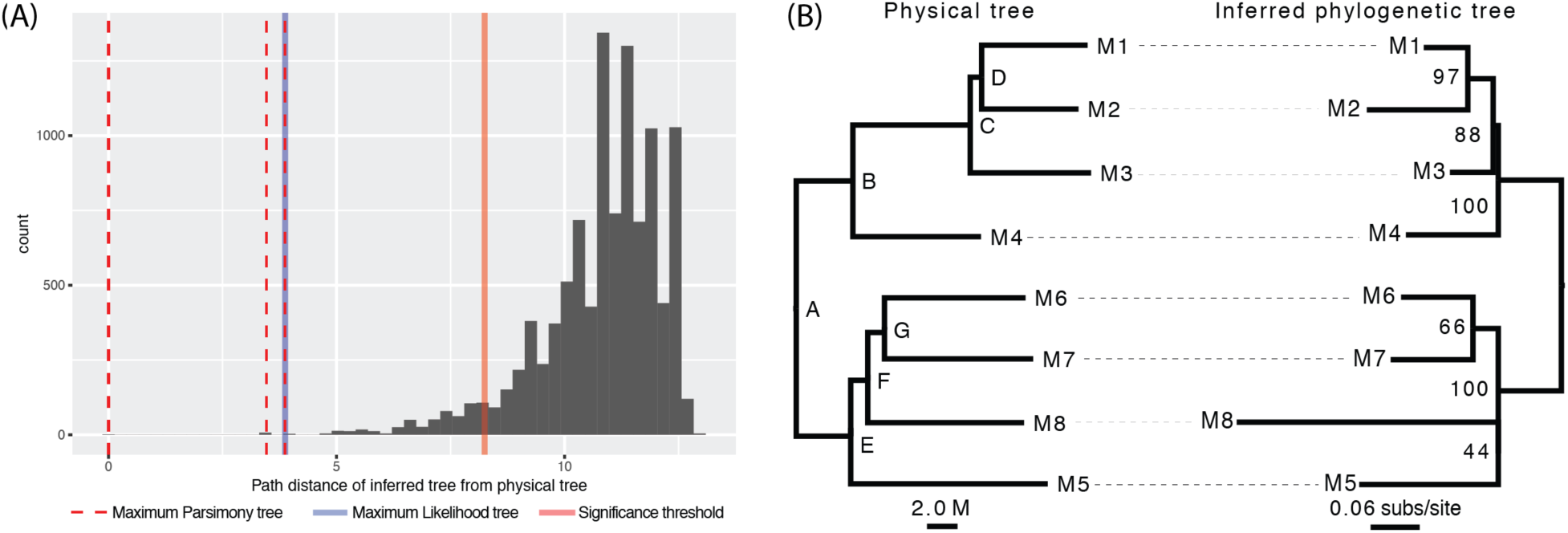
Phylogenetic trees reconstructed from somatic mutations resemble the physical structure of the tree more closely than expected by chance. **(A)** The path difference between the physical tree (Figure 1) and all 10395 possible phylogenetic trees of eight taxa is shown as a histogram. A tree with the same topology as the physical tree will have a path difference of 0. The solid red line represents the boundary of the smallest 5% of the distribution of path differences, such that a tree with a path difference lower than this line is more similar to the physical tree than expected by chance. All of the Maximum-Parsimony trees (dashed red lines) and the one Maximum-Likelihood tree (solid blue line) are more similar to the physical tree than expected by chance. **(B)** A side-by-side comparison of the physical tree (left, branch lengths in metres) and the Maximum-Likelihood tree (right, branch lengths in substitutions per site) inferred with the JC model. Letters on the nodes of the physical tree (left) correspond to the same letters of internal nodes in Figure 1. Numbers on the Maximum-Likelihood tree (right) are bootstrap percentages. There is a single difference between the two trees: the inferred tree groups samples M8 and M5 together with low bootstrap support (44%), which is a grouping that does not occur in the physical tree.

We next developed a full maximum likelihood framework to detect somatic mutations in a phylogenetic context, and used this framework to estimate the full rate and spectrum of somatic mutations in the individual *Eucalyptus melliodora* (see methods). This method increases our power to detect true somatic mutations by assuming that the phylogenetic structure of the samples follows the physical structure of the tree, an assumption that is validated by the analyses above. Using this framework, we identified 90 high-confidence somatic variants (see methods). We estimated the false negative rate by creating 14,000 in silico somatic mutations in the raw reads^23^, comprised of 1000 in silico mutations for each of the 14 branches of the physical tree, and measuring the recovery rate of these in silico mutations using our maximum likelihood approach. We were able to recover 4193 of the in silico mutations, suggesting that our recovery rate is 29.95%, and thus our false negative rate is 70.05%. Finally, we estimated our false positive rate by repeating our detection pipeline after permuting the labels of samples and replicates to remove all phylogenetic information in the data, and only considering sites that we had not previously identified as variable (see methods). By removing phylogenetic information and previously identified variable sites, we can be sure that any mutations detected by this pipeline are false positives. Across 100 such permutations, we detected 11 false-positive mutations in total, suggesting that our false positive rate is 0.11false-positive variant calls per experiment.

Based on these analyses, we can estimate the mutation rate per meter of physical growth and per year. We estimate that the true number of somatic mutations in our samples is 300 (calculated as: (90-0.11) /0.2995)). Since we sampled a total of 90.1 m of physical branch length, this equates to 3.3 somatic mutations per diploid genome per meter of branch length, or 2.75×10^−9^ somatic mutations per base per meter of physical branch length. Although the exact age of this individual is unknown, it almost certainly lies in the range 50-200 years. Given that the physical branch length connecting each sampled branch tip to the ground varies between 8.4 m and 20.3 m, we estimate that the mutation rate per base per year for a single apical meristem lies in the range 1.16×10^−10^ to 1.12×10^−9^ (i.e. 8.4 * 2.75×10^−9^ / 200 to 20.3 * 2.75×10^−9^ / 50).

With some additional assumptions it is also possible to estimate the mutation rate per generation, and to compare this to estimates from other plants. The average height of an adult *E. melliodora* individual is between 15 m and 30 m^24^, so if we assume that all somatic mutations are potentially heritable (about which there is limited evidence^1^ and ongoing discussion^25^) we can estimate that the per-generation mutation rate is between 4.13×10^−8^ and 8.25×10^−8^ mutations per base per generation. For comparison the roughly 20 cm tall *Arabidopsis thaliana* has a per-generation mutation rate of 7.1×10^−9^ mutations per base^26^. To the extent that such a comparison is accurate, which will be somewhat limited because the former estimate considers only somatic mutations and the latter considers all heritable mutations, we can then compare these estimates. Comparing the estimates suggests that despite being roughly 100 times taller than *Arabidopsis thaliana*, the per-generation mutation rate of *E. melliodora* is just ∼10 times higher, which is achieved by a roughly fifteen-fold reduction in the mutation rate per physical meter of plant growth.

Our work adds to a growing body of evidence that low somatic mutation rates per unit of growth are a general feature of many large plant species^1,2 15,16,18^. For example, a recent study of the Sitka spruce estimated a per-generation somatic mutation rate of 2.7×10^−8^, with confidence intervals that overlap ours. While this per-generation rate is high, the rate per meter of growth is very low, roughly an order of magnitude lower than our estimate for *E. melliodora*, at around 3.5×10^−10^ (2.7×10^−8^ divided by the average height of individual Sitka spruce studied of 76 m) which is strikingly similar to the rate that we estimate here^15^. Lower somatic mutation rates per unit of growth in larger plants may be the result of selection for reduced somatic mutation rates in response to the accumulation of increased genetic load in larger individuals^10,27 1, 2,15,28,29^. This pattern could also explain why larger plants tend to have lower average rates of molecular evolution than their smaller relatives^10,30^.

Several possible mechanisms might account for a reduction in accumulation of mutations per unit of growth in larger plants. Selection may favour reduction in the mutation rate per cell division through enhanced DNA repair in order to reduce the lifetime mutation risk. Alternatively, it may be that the reduction in the mutation rate is due to slower cell division. For example, plant meristems contain a slowly-dividing population of cells in the central zone of the apical meristem, and these cells are known to divide more slowly in trees than in smaller plants^31^. Indeed, the rate of cell division in the central zone is so low that one estimate put the total number of cell divisions per generation in large trees as low as one hundred^31^. Regardless of the underlying mechanism, the surprisingly low rates of somatic mutation in large plants reported here and elsewhere suggest an emerging picture in which there is a strong link between the evolution of life history and somatic mutation rates across the plant kingdom. We hope that the approach we describe here will help in further understanding this important question.

## Materials and Methods

### Data and code availability

All of the bioinformatic workflows we describe here are provided in full detail in the GitHub repository available here: https://github.com/adamjorr/somatic-variation. The raw short-read data are available online at: https://www.ncbi.nlm.nih.gov/bioproject/553104.

### Field Sampling

We used a known mosaic *Eucalyptus melliodora* (yellow box). This tree is found near Yeoval, NSW, Australia (−32.75°, 148.65°). We collected the ends of eight branches in the canopy (Figure 1). Branches were collected using an elevated platform mounted on a truck, and were placed into labelled and sealed polyethylene bags which were immediately buried in dry ice in the field. Within 24 h of collection the samples were transferred to −80°C until DNA extraction. Simultaneously, we used a thin rope to trace each branch from the tip to the main stem. These rope lengths were measured to determine the lengths of the physical branches of the tree.

### DNA Extraction, Library Preparation and Sequencing

The branches were maintained below −80°C on dry ice and in liquid nitrogen whilst sub-sampled in the laboratory. From each branch, we selected a branch tip which had at least three consecutive leaves still attached to the stem. From this branch tip we independently sub-sampled c.a. 100 mg of leaf from the ‘tip-side’ of the mid-vein on three consecutive leaves using a single hole punch into a labelled microcentrifuge tube containing two 3.5 mm tungsten carbide beads. The sealed tube was submerged in liquid nitrogen before the leaf material was ground in a Qiagen TissueLyser (Qiagen, Venlo, Netherlands) at 30 Hz in 30 s intervals before being submerged in liquid nitrogen again. This was repeated until the leaf tissue was a consistent powder, up to a total of 3.5 min grinding time.

DNA was extracted from this leaf powder using the Qiagen DNeasy Plant Mini Kit (Qiagen, Venlo, Netherlands), following the manufacturer’s instructions. DNA was eluted in 100 µl of elution buffer. DNA quality was assessed by gel electrophoresis (1% agarose in 1 × TAE containing ethidium bromide) and quantity was determined by Qubit Fluorometry (Invitrogen, California, USA) following manufacturer’s instructions.

We used a Bioruptor (Diagnode, Seraing (Ougrée), Belgium) to fragment 1µg of DNA to an average size of 300 bp (35 s on ‘High’, 30 s off for 35 cycles at 4°C). The fragmented DNA was purified using 1.6 × SeraMag Magnetic Beads (GE LifeSciences, Illinois, USA) following the manufacturer’s instructions. We used Illumina TruSeq DNA Sample Preparation kit (Illumina Inc., California, USA) following the manufacturer’s instructions to generate paired end libraries for sequencing. These libraries were sequenced on an Illumina HiSeq 2500 (Illumina Inc., California, USA) at the Biomolecular Resource Facility at the Australian National University, Canberra.

### Creation of pseudo-reference genome

Since there is no available reference genome for *E. melliodora*, we created a pseudo-reference genome by iterative mapping and consensus calling. To do this, we first mapped all of our reads to version 2.1 of the *E. grandis* reference genome ^32^using NGM ^33^, and then updated the *E. grandis* reference genome using bcftools consensus ^34^. We iteratively repeated this procedure until we saw only marginal improvement in the number of unmapped reads and reads that mapped with a mapping quality of zero. The alignment originally contained 67 M unmapped reads and 311 M reads that mapped with zero mapping quality, out of a total of 1792M reads. After the first iteration, the alignment contained 61 M unmapped reads and 349 M reads that mapped with zero mapping quality. After the last iteration, the alignment contained 59 M unmapped reads and 311 M reads that mapped with zero mapping quality. The consensus of this alignment served as the reference for all further downstream analyses.

### Variant calling for positive control

To call variants for the positive control, we mapped each replicate of each branch tip (24 samples in total) to the final pseudo-reference genome using NGM, and called genotypes using GATK 4 according to the GATK best practices workflow ^35^. This resulted in a full genome alignment of all 24 samples (three replicates of eight branches), and produced an initial set of 9,679,544 potential variable sites.

We then filtered variants to minimize the false-positive rate by retaining only those sites in which: (i) genotype calls were identical within all three replicates of each branch tip; (ii) at least one branch tip had a different genotype than the other branch tips; (iii) the site is biallelic, since multiple somatic mutations are likely to be extremely rare; (iv) the depth is less than or equal to 500, since excessive depth is a signal of alignment issues; (v) the ExcessHet annotation was less than or equal to 40, since excessive heterozygosity at a site is a sign of genotyping errors; (vi) the site is not in a repetitive region determined by a lift-over of the *E. grandis* RepeatMasker annotation, as variation in repeat regions is likely due to alignment error. This filtering produced a set of 99 high-confidence sites containing putative somatic mutations.

### Positive control

Using the set of 99 high-confidence putative somatic mutations, we use the Phangorn package in R^36^ to calculate the parsimony score of all 10,395 possible phylogenetic trees of eight taxa. This estimates the number of somatic mutations that would be required to explain each of the 10,395 phylogenetic trees, using the Fitch algorithm implemented in the Phangorn R package^36^. Of these trees, three had the maximum parsimony score of 78. One of these three trees matched the topology of the physical tree (Figure 2).

Next, we calculated the Path Difference (PD) between all 10,395 trees and the physical tree topology. The Path Difference measures differences between two phylogenetic tree topologies^37^ by comparing the differences between the path lengths of all pairs of taxa. Here we use the variant of the PD that treats all branch lengths as equal, because we are interested only in toplogical differences between trees, not branch length differences. Comparing all 10,395 trees to the physical tree topology provides a null distribution of PDs between all trees and the physical tree topology, which we can use to ask whether each of the three maximum parsimony trees is more similar to the physical tree topology than would be expected by chance. To do this, we simply ask whether the PD of each of the three observed maximum parsimony trees falls within the lower 5% of the distribution of PDs from all 10,395 trees. This was the case for all three maximum parsimony trees (p < 0.001 in all cases, see Figure 2), suggesting that our data contain biological signal which render the phylogenetic trees reconstructed from somatic mutations more similar than would be expected by chance to the physical tree.

### Variant calling for estimating the rate and spectrum of somatic mutations

Using the physical tree topology to define the relationship between samples, we called somatic mutations using DeNovoGear’s dng-call method^38^ compiled from https://github.com/denovogear/denovogear/tree/3ae70ba. Model parameters were estimated from 3-fold degenerate sites in our NGM alignment, via VCFs generated by bcftools mpileup and bcftools call with --pval-threshold=0. We estimated maximum-likelihood parameters using the Nelder-Mead numerical optimization algorithm implemented in the R package dfoptim (https://cran.r-project.org/package=dfoptim). We then called genotypes using the GATK best practices workflow as above, but with the --standard-min-confidence-threshold-for-calling argument set to 0, causing the output VCF to contain every potentially variable site in the alignment. Thus, we used GATK to generate high-quality pileups from our alignments. These pileups were then analyzed by dng-call to identify (1) heterozygous sites and (2) de novo somatic mutations. Since successful haplotype construction in a region indicates a high quality alignment, we used Whatshap 0.16 ^39^to generate haplotype blocks from the heterozygous sites.

Next, we filtered our de novo variant set to remove potential false positives. We removed variants that: (1) were on a haplotype block with a size less than 500 nucleotides; (2) were within 1000 nucleotides of another de novo variant (indicative of alignment issues); (3) had an log10 likelihood of the data (LLD) score less than −5 (indicative of poor model fit); and (4) had a de novo mutation probability (DNP) score less than 0.99999 (retaining only the highest confidence variants). This produced a final variant set of 90 variants.

### Estimation of the false negative rate

To estimate the number of mutations that we were likely to have filtered out in our variant calling pipeline, we used the method of Ness et al. ^40^, adapted to the current phylogenetic framework. Specifically, we randomly selected 14,000 sites from the first 11 scaffolds of the pseudo-reference genome, and randomly assigned 1000 of these sites to each of the 14 branches on the tree. For each of these sites, we induced in silico mutations into the raw reads with a three-step procedure. We first estimated the observed genotype at the root using DeNovoGear call at each site. We then chose a mutant genotype by mutating one of the alleles to a randomly-chosen different base using a transition/transversion ratio of 2, reflecting the observed transition/transversion ratio of eucalypts. We edited the raw reads as follows: for each mutation, we defined the samples to be mutated as all of those samples that descend from the branch on which the in silico mutation occurred. For example, an in silico mutation occurring on branch B->C in figure 1 would affect all three replicates of samples 1, 2, and 3. We then edited the reads that align to the site in question to reflect the new mutation, depending on whether the reference genotype was homozygous or heterozygous. For homozygous sites, we selected the number of reads to mutate by generating a binomially-distributed random number with a probability of 0.5 and a number of observations equal to the number of reads with the reference genotype. We then randomly selected that number of reads with the reference allele to mutate to the mutant allele, and edited the raw reads accordingly. For a heterozygous site, we edited the reads to replace all occurrences of the reference allele to mutant allele. The result of this procedure is the generation of a new set of raw fastq files, which now contain information on 1000 in silico mutations for every branch in the physical tree.

To determine the false negative rate of the variant calling pipeline, we re-ran the entire pipeline using the edited reads, and recorded the how many of the 14,000 in silico mutations were recovered by the pipeline. This number was 4193, suggesting that our false negative rates is 70.05%. In other words, we expect that our empirical analysis recovered roughly three in 10 true mutations, because our power is limited in part by attempts to filter out false positives, which also removes a number of true positives.

### Estimation of the false positive rate

To determine the false positive rate of the variant calling pipeline, we simulated random trees of our samples (where each of the eight branches is represented by three tips that denote the three replicates of that branch) by shuffling the tip labels until the tree had a maximal Robinson-Foulds distance from the original tree. This 24-taxon tree shares no splits with the original 24-taxon tree, so any phylogenetic information should be removed. We simulated 100 such trees and called variants using the pipeline above, but assuming that these trees were the physical tree, and ignoring any sites we had previously called as variable. Thus, any variants called by the pipeline must be false positives. We recovered 11 false positive calls over 100 simulations, indicating our false positive rate is approximately 0.11 calls per experiment.

### Potential functional effects

Of the 90 variants we identified, 20 were in genes. Of these, six were in coding regions, with five non-synonymous mutations and one synonymous mutation. We found no evidence that mutations were more physically clustered in the genome than would be expected by chance (p = 0.96). We detected seven mutations on the branch separating the herbivore resistant samples from the other samples. Although we lack the functional evidence to determine whether any of these mutations are directly involved in the resistance phenotype, two of the mutations occur near genes that are plausibly involved in this phenotype, and may be good candidates for further investigation. One mutation occurs near Eucgr.C00081, which is a zinc-binding CCHC type protein belonging to a small protein family known to bind RNA or ssDNA in *Arabidopsis thaliana*, and thus potentially involved in gene expression regulation. Another mutation occurs near Eucgr.I01302, an acid phosphatase that may have as a substrate phosphoenol pyruvate, and therefore may be involved in pathways associated with the production of various secondary metabolites. Intriguingly, a recent GWAS study in a related species, *Eucalyptus polybractea*, found a very strong association between variants in the *ppt2* gene (phosphoenol pyruvate transporter 2) and the total foliar concentration of sesquiterpenes, key secondary metabolites known to be involved in herbivore defence^41^.

## References

1. Plomion, C. et al. Oak genome reveals facets of long lifespan. Nat Plants 4, 440–452 (2018).

2. Schmid-Siegert, E. et al. Low number of fixed somatic mutations in a long-lived oak tree. Nat Plants 3, 926–929 (2017).

3. Whitham, T. G. & Slobodchikoff, C. N. Evolution by Individuals, Plant-Herbivore Interactions, and Mosaics of Genetic-Variability – the Adaptive Significance of Somatic Mutations in Plants. Oecologia 49, 287–292 (1981).

4. Buss, L. W. Evolution, Development, and the Units of Selection. P Natl Acad Sci USA 80, 1387–1391 (1983).

5. Walbot, V. On the life strategies of plants and animals. Trends Genet 1, 165–169 (1985).

6. Sutherland, W. J. & Watkinson, A. R. Somatic Mutation – Do Plants Evolve Differently. Nature 320, 305–305 (1986).

7. Klekowski, E. J. & Godfrey, P. J. Ageing and mutation in plants. Nature 3, 389–391 (1989).

8. Ally, D., Ritland, K. & Otto, S. P. Aging in a long-lived clonal tree. PLoS Biol 8, (2010).

9. Gill, D. E., Chao, L., Perkins, S. L. & Wolf, J. B. Genetic Mosaicism in Plants and Clonal Animals. Annual Review of Ecology and Systematics 26, 423–444 (1995).

10. Lanfear, R. et al. Taller plants have lower rates of molecular evolution. Nat Commun 4, 1879 (2013).

11. Khoury, C., Laliberté, B. & Guarino, L. Trends in ex situ conservation of plant genetic resources: a review of global crop and regional conservation strategies. Genet Resour Crop Evol 57, 625–639 (2010).

12. Schoen, D. J., David, J. L. & Bataillon, T. M. Deleterious mutation accumulation and the regeneration of genetic resources. P Natl Acad Sci USA 95, 394–399 (1998).

13. Michel, A. et al. Somatic mutation-mediated evolution of herbicide resistance in the nonindigenous invasive plant hydrilla (Hydrilla verticillata). Mol Ecol 13, 3229–3237 (2004).

14. Klekowski, E. J., Jr., Corredor, J. E., Morell, J. M. & Del Castillo, C. A. Petroleum pollution and mutation in mangroves. Marine Pollution Bulletin 28, 166–169 (1994).

15. Hanlon, V. C. T., Otto, S. P. & Aitken, S. N. Somatic mutations substantially increase the per-generation mutation rate in the conifer Picea sitchensis. Evolution Letters (2019). doi:10.1002/evl3.121

16. Wang, L. et al. The architecture of intra-organism mutation rate variation in plants. PLoS Biol 17, e3000191 (2019).

17. Schultz, S. T. & Scofield, D. G. Mutation accumulation in real branches: fitness assays for genomic deleterious mutation rate and effect in large-statured plants. Am Nat 174, 163–175 (2009).

18. Bobiwash, K., Schultz, S. T. & Schoen, D. J. Somatic deleterious mutation rate in a woody plant: estimation from phenotypic data. Heredity 111, 338–344 (2013).

19. Padovan, A., Lanfear, R., Keszei, A., Foley, W. J. & Külheim, C. Differences in gene expression within a striking phenotypic mosaic Eucalyptus tree that varies in susceptibility to herbivory. BMC Plant Biol 13, 29 (2013).

20. Edwards, P. B., Wanjura, W. J., Brown, W. V. & Dearn, J. M. Mosaic resistance in plants. Nature 347, 434–434 (1990).

21. Grattapaglia, D. et al. Progress in Myrtaceae genetics and genomics: Eucalyptus as the pivotal genus. Tree Genetics & Genomes 8, 463–508 (2012).

22. McKenna, A. et al. The Genome Analysis Toolkit: a MapReduce framework for analyzing next-generation DNA sequencing data. Genome Res 20, 1297–1303 (2010).

23. Keightley, P. D., Ness, R. W., Halligan, D. L. & Haddrill, P. R. Estimation of the Spontaneous Mutation Rate per Nucleotide Site in a Drosophila melanogaster Full-Sib Family. Genetics 196, 313–320 (2014).

24. Boland, D. J.et al. Forest trees of Australia. (CSIRO, 2006).

25. Lanfear, R. Do plants have a segregated germline? PLoS Biol 16, e2005439 (2018).

26. Ossowski, S. et al. The Rate and Molecular Spectrum of Spontaneous Mutations in Arabidopsis thaliana. Science 327, 92–94 (2010).

27. Zhong, B., Fong, R., Collins, L. J., McLenachan, P. A. & Penny, D. Two New Fern Chloroplasts and Decelerated Evolution Linked to the Long Generation Time in Tree Ferns. Genome Biology and Evolution 6, 1166–1173 (2014).

28. Klekowski, E. J. Mutation, Developmenal Selection, and Evolution. (Columbia University Press, 1988).

29. Scofield, D. G. A definitive demonstration of fitness effects due to somatic mutation in a plant. 112, 361–362 (2014).

30. Smith, S. A. & Donoghue, M. J. Rates of molecular evolution are linked to life history in flowering plants. Science 322, 86–89 (2008).

31. Romberger, J. A., Hejnowicz, Z. & Hill, J. F. Plant Structure: Function and Development. (Springer, 1993).

32. Bartholomé, J. et al. High-resolution genetic maps of Eucalyptus improve Eucalyptus grandis genome assembly. New Phytol 206, 1283–1296 (2014).

33. Sedlazeck, F. J., Rescheneder, P. & Haeseler von, A. NextGenMap: fast and accurate read mapping in highly polymorphic genomes. Bioinformatics 29, 2790–2791 (2013).

34. Li, H. A statistical framework for SNP calling, mutation discovery, association mapping and population genetical parameter estimation from sequencing data. Bioinformatics 27, 2987–2993 (2011).

35. Van der Auwera, G. A. et al. From FastQ Data to High-Confidence Variant Calls: The Genome Analysis Toolkit Best Practices Pipeline. Current Protocols in Bioinformatics 43, 11.10.1–11.10.33 (2013).

36. Schliep, K. P. phangorn: phylogenetic analysis in R. Bioinformatics 27, 592–593 (2011).

37. Steel, M. A. & Penny, D. Distributions of tree comparison metrics-some new results. Systematic Biology 42, 126–141 (1993).

38. Ramu, A. et al. DeNovoGear: de novo indel and point mutation discovery and phasing. Nat. Methods 10, 985–987 (2013).

39. Martin, M. et al. WhatsHap: fast and accurate read-based phasing. bioRxiv 085050 (2016). doi:10.1101/085050

40. Ness, R. W., Morgan, A. D., Colegrave, N. & Keightley, P. D. Estimate of the spontaneous mutation rate in Chlamydomonas reinhardtii. Genetics 192, 1447–1454 (2012).

41. Kainer, D. et al. High marker density GWAS provides novel insights into the genomic architecture of terpene oil yield in Eucalyptus. New Phytol Online early (2019).

